# Peroxidasin protein expression and enzymatic activity in metastatic melanoma cell lines are associated with invasive potential

**DOI:** 10.1101/2021.06.04.447036

**Authors:** Martina Paumann-Page, Nikolaus F. Kienzl, Jyoti Motwani, Boushra Bathish, Louise N. Paton, Nick J. Magon, Benjamin Sevcnikar, Paul G. Furtmüller, Michael W. Traxlmayr, Christian Obinger, Mike R. Eccles, Christine C. Winterbourn

## Abstract

Peroxidasin, a heme peroxidase, has been shown to play a role in cancer progression. mRNA expression has been reported to be upregulated in metastatic melanoma cell lines and connected to the invasive phenotype, but little is known about how peroxidasin acts in cancer cells. We have analyzed peroxidasin protein expression and activity in eight metastatic melanoma cell lines using an ELISA developed with an in-house peroxidasin binding protein. RNAseq data analysis confirmed high peroxidasin mRNA expression in the five cell lines classified as invasive and low expression in the three non-invasive cell lines. Protein levels of peroxidasin were higher in the cell lines with an invasive phenotype. Active peroxidasin was secreted to the cell culture medium, where it accumulated over time, and peroxidasin protein levels in the medium were also much higher in invasive than non-invasive cell lines. The only well-established physiological role of peroxidasin is in the formation of a sulfilimine bond, which cross-links collagen IV in basement membranes via catalyzed oxidation of bromide to hypobromous acid. We found that peroxidasin secreted from melanoma cells formed sulfilimine bonds in uncross-linked collagen IV, confirming peroxidasin activity and hypobromous acid formation. Moreover, 3-bromotyrosine, a stable product of hypobromous acid reacting with tyrosine residues, was detected in invasive melanoma cells, substantiating that their expression of peroxidasin generates hypobromous acid, and showing that it does not exclusively react with collagen IV, but also with other biomolecules.

## INTRODUCTION

Invasion and metastasis are fundamental hallmarks of tumor biology and the main causes of cancer mortality. There is mounting evidence that expression of the heme peroxidase, peroxidasin (PXDN), promotes cancer progression and the invasion of tumor cells. The first evidence of PXDN expression in melanoma was reported in 1994 by Weiler et al., where PXDN was termed melanoma-associated antigen MG50 (1). A more recent study showed that PXDN was vastly over-expressed in metastatic melanoma cell lines with an invasive phenotype (2). Moreover, *PXDN* gene silencing led to reduced melanoma cell invasion *in vitro* as well as *in vivo*. High expression of PXDN has also been reported for several other cancers (3, 4) including ovarian (5), prostate (6, 7), breast (8, 9) and brain cancer (10). For ovarian, breast and prostate cancer, a correlation with poor prognosis was found (5, 9) (6), which was also reported for endometrial, cervical and stomach cancer (4). However, the mechanistic role of PXDN in cancer remains mostly unexplored and may depend on the type of cancer as well as on the stage.

PXDN, a member of the peroxidase-cyclooxygenase superfamily (11, 12), is a homotrimeric multidomain peroxidase which is secreted to the extracellular matrix (ECM). PXDN mRNA is expressed in most tissues and many cell types (13, 14). The enzyme is known to play an important role in the stabilization of basement membranes (BM), which is a specialized form of ECM, by forming a covalent cross link in collagen IV, one of the main constituents of BMs (15). This cross link is crucial for BM integrity and contributes to tensile strength and stiffness of the three-dimensional supramolecular structures formed by collagen IV and other basement membrane proteins (16). PXDN reacts with hydrogen peroxide to oxidize bromide ions to hypobromous acid (HOBr), which subsequently forms the sulfilimine bond between a methionine and a hydroxylysine residue of two juxtaposed non-collagenous domains in collagen IV (15). In addition to bromide PXDN effectively oxidizes iodide and thiocyanate, leading to the formation of hypoiodous acid and hypothiocyanite, respectively (17). PXDN also reacts with a wide range of other typical peroxidase substrates including ascorbate, tyrosine, serotonin, nitrite, urate and (poly)phenolic compounds, resulting in the formation of free radical species (18). Depending on hydrogen peroxide levels and the availability of substrates, all these reactions may occur intracellularly and in the extracellular space after secretion of PXDN.

HOBr as well as other oxidants and secondary reaction products (e.g. bromamines) have the capacity to lead to numerous intra- and extracellular posttranslational oxidative modifications of biomolecules including BM and ECM proteins, cell surface receptors and signaling cascades (19). Conformational changes of modified proteins may alter the susceptibility or resistance to proteolytic degradation and oxidants may furthermore impact on DNA and RNA integrity, function, stability and DNA methylation (20–23). Moreover, activity of PXDN was shown to promote angiogenesis via ERK1/2, AKT and FAK pathways (24) and is required for ECM-mediated signaling via ERK1/2 and FAK, influencing cell growth and survival (25). This highlights how PXDN activity has the potential to influence many aspects of tumor biology.

PXDN may also act as a signaling protein. In addition to its catalytically active peroxidase domain, PXDN possesses an N-terminal leucine-rich repeat domain and four immunoglobulin domains as well as a C-terminal von Willebrand factor C module (12, 26). These domains are typical for the ECM and known for participating in protein-protein and protein-cell surface interactions. Direct binding of PXDN to laminin has been observed (27). The ECM influences all stages of cancer development and progression. It is an important platform for biochemical and biophysical signals that influence cell functions including adhesion, differentiation, proliferation, survival, migration, invasion and angiogenesis (28, 29). In order to understand whether PXDN contributes to melanoma progression by affecting any of these activities, it is necessary to know more about its protein expression in relation to invasiveness of melanoma cells, whether it is active, and the reactions it undergoes. To address these questions, we have measured PXDN protein expression and activity in invasive and non-invasive metastatic melanoma cell lines from the well characterized New Zealand melanoma (NZM) panel. We examined if mRNA expression is associated with invasive phenotype and used an ELISA with an in-house developed PXDN binding protein to show that mRNA levels translate to protein expression levels. Using a peroxidase assay and detection of 3-bromotyrosine we show that PXDN is active and forms HOBr in high expressing cells. We furthermore demonstrate secretion of active PXDN which is capable of forming sulfilimine bonds in uncross-linked collagen IV.

## MATERIAL AND METHODS

### Reagents

All reagents used were of the highest available analytical quality and were from Sigma Aldrich unless stated otherwise. The concentration of hydrogen peroxide was determined at 240 nm using an extinction coefficient of 43.6 M^-1^cm^-1^. Human myeloperoxidase was from Planta (Vienna, Austria) and lactoperoxidase (L2005) was from Sigma-Aldrich. Formic acid ≥98 % was purchased from GPR RECTAPUR® (VWR International, Singapore), water for chromatography (LC-MS Grade) and acetonitrile hypergrade for LC-MS was from LiChrosolv® (Merck KGaA, Darmstadt, Germany).

### Culture of metastatic melanoma cells

The eight New Zealand melanoma cell lines (NZM6, NZM9, NZM11, NZM12, NZM15, NZM22, NZM40, NZM42) were originally generated from surgical samples of human metastatic melanoma tumors as previously described, after written consent was obtained from all patients under Auckland Area Health Board Ethics Committee guidelines (30). The cell lines were authenticated and generously provided by B. Baguley from the University of Auckland, Auckland, New Zealand. Melanoma cell lines were grown in MEMα (Minimum Essential Medium α, no nucleosides; 12561056, Gibco™) supplemented with 5 % fetal bovine serum, insulin (5 μg/mL), transferrin (5 μg/mL) and sodium selenite (5 ng/mL) (Roche Diagnostics, Germany), 100 units/mL of penicillin, 100 μg/mL of streptomycin. A normal human epidermal neonatal lightly pigmented melanocyte cell line (HEMn-LP) was used as a reference cell line. HEMn-LP was cultured in Medium 254 (M254500, Gibco™) supplemented with 1 % human melanocyte growth supplement-2 (0165, Gibco™). All cell lines were cultured in a humidified 37 °C hypoxic cell culture incubator (Xvivo System Model X3, BioSpherix) at 5 % O_2_, 5 % CO_2_ and 90 % nitrogen, unless stated otherwise. Low oxygen tension was used in order to mimic physiologically low oxygen levels in tumors. All cell lines were used at low passage numbers (< 30) and sub-cultured when confluence reached 70 – 90 %. All cell lines tested negative for mycoplasma (e-Myco™ VALiD Mycoplasma PCR Detection Kit).

### Cell growth, lysis and determination of protein concentration

Cells were routinely grown in 100 mm cell culture dishes or 6 well plates (3-4 million cells per 100 mm plate at full confluence, depending on cell line characteristics). Cell lysates and cell culture medium were analyzed for PXDN. For PXDN to accumulate in the medium, cells were grown to full confluence followed by a medium exchange and cells and medium were harvested three days later. Cells were harvested after taking off the medium and two washes with phosphate buffered-saline (PBS). Cells were subsequently lysed on ice by adding 200 μL of lysis buffer (10 mM Tris pH 8, 1 mM EDTA, 1 % w/v sodium deoxycholate, 1x Complete® Protease Inhibitor) and plates were scraped with a cell scraper. Cell lysates were sonicated on ice with a fine tip probe three to four times for ten seconds each, until lysates were homogeneous. Secreted PXDN in the cell medium was analyzed after any detached cells were removed by centrifugation at 500 g for 10 min. A Direct Detect® Infrared Spectrometer was used to measure the protein concentration of cell lysates, which were typically between 5 – 10 mg/mL for different cell lines. The protein concentration of the cell culture medium was typically ∼2 mg/mL, predominantly originating from 5 % fetal bovine serum added to the medium.

### Recombinant PXDN

Recombinant PXDN consisting of the four immunoglobulin domains and the peroxidase domain has been extensively characterized previously and was termed as hsPxd01-con4, being one of several truncated constructs in previous works (17, 18, 26, 31). In this manuscript hsPxd01-con4 will be simply referred to as PXDN-con4. PXDN-con4 has an N-terminal His-tag. The concentration of PXDN-con4 was determined using the extinction coefficient of 147,500 M^-1^ cm^-1^ per heme and conversions to micrograms were made using a molar mass of 140 kDa.

### Development of PXDN-specific binding proteins based on rcSso7d using yeast display

Binders based on rcSso7d (reduced charge Sso7d, which is a charge-neutralized version of the 7 kDa protein Sso7d from hyperthermophilic archaeon *Sulfolobus solfataricus* with charge-neutralized binding site) that specifically recognize human PXDN were engineered using yeast surface display technology, according to previously published protocols (32–34). Briefly, the yeast-displayed libraries rcSso7d-11 and rcSso7d-18 were selected for binding to biotinylated PXDN-con4 (PXDN-con4-biotin). Biotinylation was carried out using EZ-Link Sulfo-NHS-LC Biotin, Thermo Fisher. Alternatively, to avoid enrichment of binders recognizing biotinylated epitopes on PXDN-con4-biotin, non-biotinylated PXDN-con4 was used in some sorting rounds, as described below. All yeast display selections and titrations were performed in PBS supplemented with 0.1 % bovine serum albumin. Binders expressed on yeast surface contained two tags for immunofluorescent detection (a hemagglutinin (HA) epitope tag at the N-terminus and a c-myc epitope tag at the C-terminus).

The yeast display selection process was started with two rounds of selections using streptavidin-coated dynabeads (Thermo Fisher) loaded with PXDN-con4-biotin, followed by error prone PCR (epPCR) using 2 µM each of 8-oxo-2’
s-deoxyguanosine 5’-triphosphate (8-oxo-dGTP) and 2’-deoxy-P-nucleoside 5’-triphosphate (dPTP) (32). After an additional round of magnetic bead selection, the libraries were further enriched by four rounds of flow cytometric sorting. Labeling of the cells for flow cytometry involved a primary incubation with PXDN-con4-biotin (concentrations between 100 and 200 nM, depending on the sorting round) and 5 µg/mL mouse anti-c-myc (clone 9E10, Thermo Fisher), followed by a secondary incubation with 20 µg/mL goat anti-mouse IgG-Alexa Fluor 488 and 10 µg/mL streptavidin-Alexa Fluor 647 (both Thermo Fisher). In sorting rounds 3 and 4, the cells were incubated with 100 nM non-biotinylated PXDN-con4, followed by a secondary incubation with 5 µg/mL anti-HA-Alexa Fluor 647 (clone 16B12, BioLegend) and 5 µg/mL anti-Penta-His-Alexa Fluor 488 (Qiagen), which recognized the His-tag on PXDN-con4. After four flow cytometric sorting rounds, enriched PXDN-con4-specific rcSso7d-based binders were sequenced, yielding four different sequence families defined as a group of binders with similar sequence patterns in their engineered binding sites and likely recognizing the same epitope.

To further improve their affinities, four epPCR libraries were constructed based on binders (POX1, POX3, POX6 and POX9) derived from those four different sequence families. These libraries were selected for PXDN-con4 binding in two rounds of flow cytometric sorting, followed by another epPCR and finally three rounds of flow sorting. To push the selection pressure towards high affinity, the antigen concentration was steadily decreased with progressing sorting rounds from 50 to 2 nM. Finally, plasmids encoding the selected binders were isolated from the yeast cells, used for *E. coli* transformation and sequenced.

To analyze the affinities of enriched binders, the yeast strain EBY100 (35) was transformed with plasmids encoding individual binders. After expression on the yeast surface, individual mutants were tested for binding to soluble non-biotinylated PXDN-con4, with PXDN-con4 concentrations ranging from 0.8 to 600 nM. PXDN-con4-binding intensities were analyzed by calculating the median fluorescence intensity (MFI) of the displaying population (i.e., HA-positive cells). Inter-experimental variation of the maximal MFI values (due to, e.g., different yeast surface expression levels between different experiments, which directly result in different maximal antigen binding levels) was compensated by multiplying all data points of a given experiment with a constant. This constant was chosen such that the average of the MFIs of all mutants at 600 nM is identical between different experiments. Thus, the shape of the curves, as well as the MFI ratios between the different mutants were not changed but experiments with lower overall intensities were not under-represented. A 1:1 binding model was fitted to the data. Competition experiments were performed to determine overlapping epitopes between different binders (supplementary Figure 2E).

### Soluble expression of enriched, high affinity PXDN-binders

Enriched PXDN-binders were sub-cloned into the pE-SUMO vector (Life-Sensors) and expressed in *E. coli* as fusion proteins comprising an N-terminal His_6_-tag, followed by small ubiquitin like-modifier (SUMO) and the respective binder, as described previously (34). After purification of the His_6_-SUMO-binder fusion proteins using TALON metal affinity resin (Clontech), they were digested with SUMO protease 1, resulting in cleavage just before the N-terminus of the rcSso7d mutants. The digested products were again purified using TALON resin and the cleaved binders were collected from the flow through. A detailed description of the expression and purification of rcSso7d-based binders was published previously (34). Plasmids encoding all engineered PXDN-specific binders will be made available to others.

### Surface plasmon resonance (SPR)

The affinity between the engineered POX10 binder (an affinity-matured derivative of POX9) and PXDN-con4 was additionally analyzed by SPR using a Biacore T200 instrument (GE Healthcare). A superfolder green fluorescent protein POX10 fusion protein (sfGFP-POX10) was used for this experiment, which was also sub-cloned into a pE-SUMO vector and expressed with an N-terminal His_6_-tag, the POX10 binder sequence, two G4S linkers (four glycine and one serine residue) and the superfolder GFP protein. sfGFP-POX10 was diluted to 5 µg/mL in sodium acetate buffer (pH 4.5) and subsequently covalently immobilized on a CM5 chip by amine coupling according to the manufacturer’
ss protocol (GE Healthcare). Next, PXDN-con4 was diluted to concentrations ranging from 3.125 to 200 nM in PBS with 0.1 % bovine serum albumin buffer containing 0.05 % Tween-20. These samples, as well as buffer only controls, were applied to the sfGFP-POX10-coated chip for 450 s (flow rate of 30 µL/min), followed by injection of buffer only. After buffer baseline subtraction, data were fitted using a kinetic 1:1 binding model using the Biacore T200 Evaluation Software (GE Healthcare).

### Peroxidasin ELISA

We developed an indirect sandwich ELISA using the His_6_-SUMO-POX10 fusion protein (referred to as PXDN binding protein) to capture PXDN and subsequently measure PXDN activity and PXDN protein in one assay. Corning high binding ELISA microplates were incubated with the PXDN binding protein (200 ng/well in 50 μL of PBS) over night at room temperature, followed by two hours of blocking with 75 μL/well of assay buffer (1 % BSA, 0.025 % Tween-20 in PBS). Between all ELISA steps plates were washed three times with PBS. Samples (cell lysates or cell culture medium) were applied (50 μL/well) and incubated for 2 hours at 37 °C. Recombinant PXDN-con4 was routinely applied to each plate and used to create a standard curve for PXDN activity (3.125 – 200 ng/well in 50 μL assay buffer). PXDN activity was measured by adding 50 μL/well of 50 μM Amplex™ UltraRed Reagent (Thermo Fisher Scientific), 20 μM hydrogen peroxide and 50 mM bromide in 50 mM phosphate buffer pH 7.4. The presence of bromide increased the relative fluorescence by approximately factor 2. The plate was incubated for exactly 30 min in the dark at 37 °C before fluorescence of the formed resorufin was measured (λ_ex_ 544 nm, λ_em_ 590 nm). An in-house produced rabbit anti-PXDN antiserum (raised against the C-terminal peptide consisting of amino acids 1329 to 1479; antigen was kindly provided by M. Geizst, Budapest; (14)) was added as primary detection antibody (1:1000), followed by a biotinylated goat anti-rabbit antibody (1:2000) and an Extra-Avidin alkaline phosphatase antibody (1:1000), diluted in assay buffer, 50 μL/well, 1 h incubation at 37 °C. 4-nitrophenyl phosphate disodium salt hexahydrate (pNPP) in 10 % diethanolamine containing 0.5 mM MgCl_2_ was added (2 mg/mL, 50 μL/well) and absorbance was measured at 405 nm after incubating at 37 °C for exactly 60 minutes (to allow comparability between assays). Standards were analyzed in duplicate and samples were analyzed in duplicate or triplicate.

### MS method for detection of PXDN

A tryptic digest of 4 μg of PXDN-con4 was used to identify two peptides (AGEIFER and AFFSPFR) for PXDN detection by LC-MS/MS in complex samples. In-gel tryptic digestion was performed on 400 μg total protein cell lysates and medium samples in order to remove detergents from the lysis buffer and to reduce background noise and ion suppression (36). Briefly, to immobilize proteins, sample volumes were reduced to 50 μL and mixed with 50 μL of 40 % w/v acrylamide and polymerized by adding 5 μL N,N,N’,N’-tetramethyl-ethylenediamine and 5 μL of 10 % w/v ammonium persulfate (APS). The gel was cut into 1 mm^3^ pieces, fixed, reduced and alkylated. In gel-digest was performed using 20 μg of trypsin in 1.5 mL of 10 mM ammonium bicarbonate (ABC) buffer with 10 % v/v acetonitrile according to the reference. After extraction the supernatant was dried in a speed vac and samples were reconstituted in 20 μL of 0.1 % v/v formic acid. A tryptic digest of PXDN-con4 was used to semi-quantitatively estimate PXDN content of the samples.

A multiple reaction monitoring LC-MS/MS method was used with chromatographic separation on an Agilent Technologies 1290 Infinity LC system (Santa Clara, CA, USA). The autosampler was set to 4 °C and the column compartment to 40 °C. The column was a Jupiter® 4 µm Proteo 90 Å, LC Column, 150 x 2 mm from Phenomenex® (Torrance, CA, USA), operated at a flow rate of 0.25 mL/min. A sample volume of 3 μL was injected. Mobile phases used were 0.1 % v/v formic acid in water (A) and 0.1 % v/v formic acid in acetonitrile (B). After an initial isocratic step for 5 minutes at 2 % B a linear gradient was run to 30 % B over 20 minutes followed by a step to 98 % B before re-equilibration at 2 % B.

Mass spectrometry was performed with a QTRAP®6500 (AB SCIEX, Framingham, MA, USA) equipped with an IonDrive™ Turbo V Ion Source operated in positive ionization mode. Analyst® 1.6.3 (AB SCIEX, Framingham, MA, USA) was used for data acquisition. The optimized acquisition parameters were as follows: curtain gas: 15, ionspray voltage: 5500, temperature: 400, ion source gas 1: 35, ion source gas 2: 35, declustering potential: 90, exit potential: 10, collision cell potential: 14. Dwell time was 100 ms and the collision gas was nitrogen. Data were analyzed using PeakView® 2.2 (AB SCIEX, Framingham, MA, USA) to determine the area under the curve for monitored transitions. The transitions monitored for PXDN are in Supplementary Table 1. AGEIFER y3 and AFFSPFR y5 gave the strongest signals and were used for quantification.

### Cross-linking of collagen IV by peroxidasin secreted from melanoma cell lines

Mouse epithelial PFHR9 PXDN wild type cells (WT) and PXDN knock out (KO; kindly provided by Dr Jay Bhave) (37) were grown in DMEM containing 10 % FBS and 1 % penicillin/streptomycin antibiotics at 37 °C under humidified 5 % CO_2_, as previously reported (38). Cells were seeded in 10 cm cell culture plates at approximately 70 % confluence and grown for eight to ten days to deposit ECM before hypotonic lysis to remove cells (39) to isolate WT and KO ECM. Melanoma cell lines were grown in 100 mm plates, trypsinized at 80 - 100 % confluence, seeded on top of the ECM isolated from PFHR9 KO cells and incubated at 5 % CO_2_ at ambient oxygen concentration. To inhibit PXDN and subsequently the formation of the crosslink, 100 μM phloroglucinol (PHG; 1,3,5-trihydroxybenzene) was added to the medium when cells were seeded and 24 hours later, where indicated. After 48 h the medium was removed and cells were washed twice with PBS. Cells and ECM were lysed on ice with 1 mL of lysis buffer (10 mM Tris pH 8, 1 mM EDTA, 1 % w/v sodium deoxycholate, 1x Complete® Protease Inhibitor) and plates were scraped. Lysates were sonicated on ice, spun at 20,000 g for 20 min at 4 °C and the supernatant was discarded. The ECM-containing pellet was washed twice in wash buffer (1 M NaCl, 10 mM Tris pH 7.5, 20,000 g for 10 min at 4 °C). For the isolation of the non-collagenous domain (NCD) the ECM was digested with collagenase (38). NCD of samples were resolved by SDS-PAGE on 12 % gels under non-reducing conditions and stained with Coomassie Brilliant Blue R-250. Monomer and dimer bands of the NCD were analyzed by densitometry (Alliance Uvitec Cambridge imager).

### 3-Bromotyrosine detection by LC/MS/MS

Melanoma cell lines NZM40 and NZM42 were seeded at 70 % confluence in 6 well plates. After 18 h the medium was exchanged to MEMα without phenol red (41061029, Gibco™) and bromide (100 and 200 μM, respectively) or phloroglucinol (100 μM) were added daily after medium exchange for 5 days before cells were harvested. Bromotyrosine detection and quantification was performed using a stable isotope dilution LC/MS/MS method as recently described by Bathish et al (38) based on (40) and (41).

## RESULTS

To substantiate previous findings of high *PXDN* mRNA expression in invasive melanoma cells (2) and establish whether this is associated with high expression of active PXDN protein, we examined eight cell lines from the New Zealand melanoma (NZM) cell line collection, which were originally generated from surgical samples of human metastatic melanoma tumors (30) (42). As previously reported (43, 44), five of the cell lines were classified as invasive (NZM6, NZM9, NZM11, NZM22, NZM40) and three as non-invasive (NZM12, NZM15, NZM42) using *in-vitro* Matrigel and Boyden chamber invasion assays. These analyses showed at least a 20-fold difference in invasion behavior, with no overlap between the groups.

### PXDN RNAseq analysis

Analysis of the RNAseq data reported by Motwani et al. (43) (GSE153592) showed that mRNA expression of *PXDN* was highly elevated in each of the invasive metastatic melanoma cell lines compared to the non-invasive cell lines (Figure 1). Expression levels varied within the invasive cell lines with NZM40 expressing vastly elevated *PXDN* mRNA. The mean value of the group of invasive cell lines is approximately 100 times higher than in the non-invasive group. Variable expression was confirmed in a larger cohort of 40 NZM cell lines (not assessed for invasion), 18 of which showed no detectable *PXDN* mRNA expression and the others ranging from low levels up to that seen with NZM40. None of the cell lines investigated express any other peroxidases as determined from the RNAseq data (myeloperoxidase, lactoperoxidase, eosinophil peroxidase and thyroid peroxidase).

**Figure 1:**
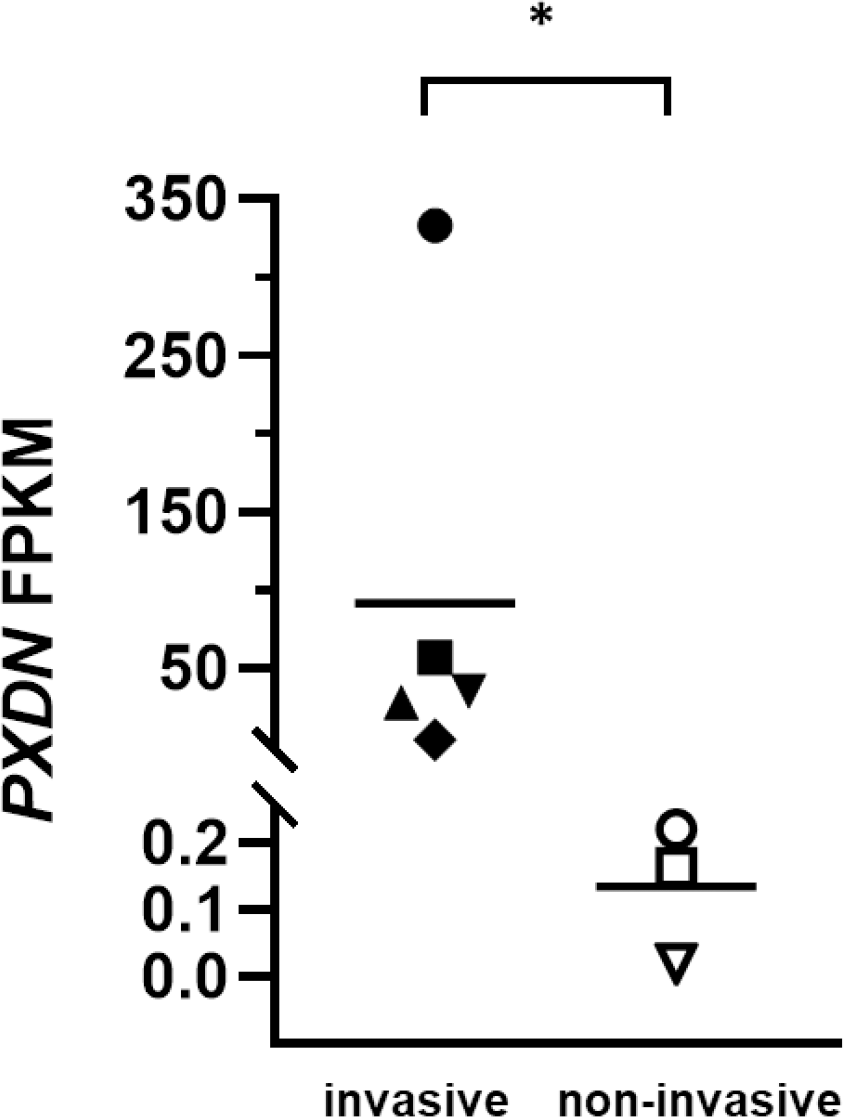
*PXDN* mRNA expression of melanoma cell lines. FPKM (Fragments per kilobase of transcript per million mapped reads) values of *PXDN* mRNA from RNAseq of melanoma cell lines grouped by their invasive potential extracted from RNAseq data obtained by Motwani et al (43). Invasive (▲) NZM6, (♦) NZM9, (◼) NZM11, (▼) NZM22, (•) NZM40 and non-invasive cell lines (○) NZM42, (□) NZM12, (▵) NZM15; means and individual points, *, *P* =0.036 by two-tailed Mann Whitney test.

### Generation of PXDN-specific binding proteins

In order to examine the melanoma cells for PXDN protein expression and activity, it was necessary to develop sensitive assays. Since the availability of effective anti-PXDN antibodies is limited, so we used a synthetic approach to developing high affinity PXDN-specific binding proteins. As a protein scaffold, we chose a recently introduced charge-neutralized variant of the hyperthermostable protein Sso7d (termed reduced charge Sso7d, rcSso7d), which was subsequently engineered for specific PXDN recognition using yeast surface display (32–34). Briefly, nine surface exposed residues were randomly mutated (cyan residues in Figure 2A) and the resulting rcSso7d-libraries were displayed on the surface of yeast and selected for binding to PXDN-con4. After several selection rounds, representative binders from four different families, POX1, POX3, POX6 and POX9, were selected (sequences shown in supplementary Figure 1). They and their affinity-matured descendants were displayed on the surface of yeast and their binding affinities with PXDN-con4 were assessed. As shown for the most efficient mutant in Figure 2B and the others in supplementary Figure S2A-C, compared with their parental variants (black curves), all affinity-matured variants (blue or red curves) showed improved affinities down to the low nanomolar range, which is comparable to those typically obtained with antibodies. Expression of these engineered binders in *E. coli* yielded monomeric, non-aggregating proteins, as shown by size exclusion chromatography (SEC; Supplementary Figure S2D). POX5 was non-competitive with POX11, which is a close relative of POX10 (Supplementary Figure S2E). This suggests that POX11 (or POX10) and POX5 bind to different epitopes on PXDN. We chose POX10 as the most promising PXDN-binder, as it combines a high affinity to PXDN-con4 with low retention on the SEC column (suggesting no or low non-specific interaction). In its soluble form expressed in *E. coli* POX10 bound to PXDN-con4 with a *K*_d_ = 74 ± 5 nM as demonstrated by surface plasmon resonance (Figure 2C).

**Figure 2:**
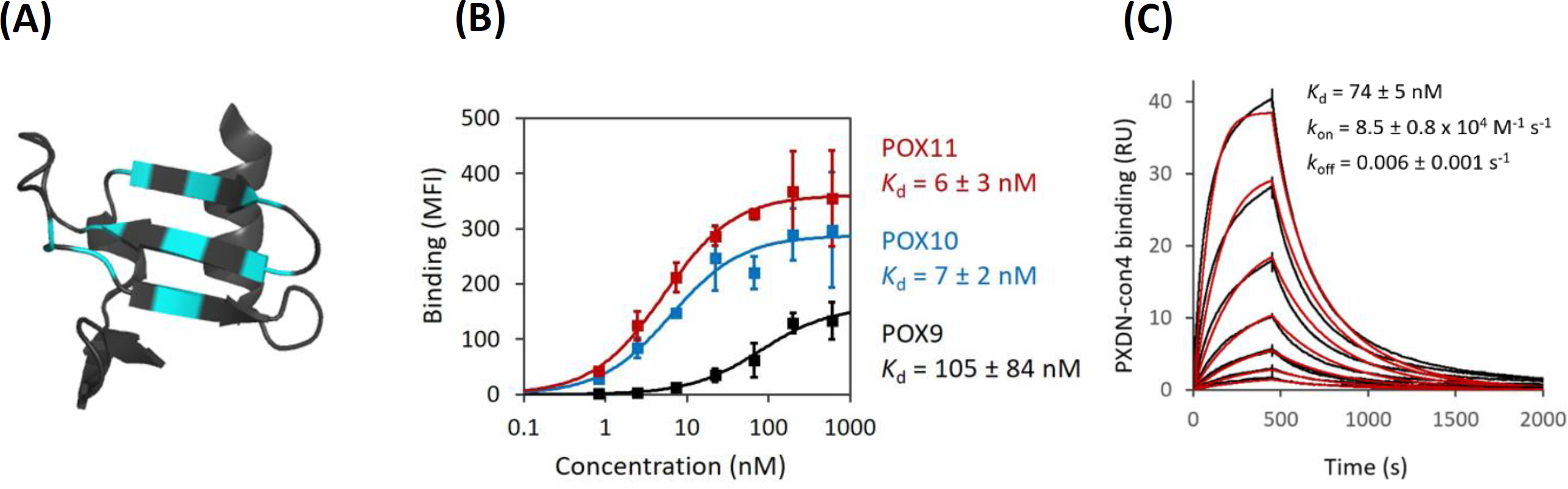
Engineering of PXDN-specific binders. **(A)** Crystal structure of the rcSso7d binding scaffold. Depicted is a hRBP4/A1120-specific mutant (PDB-ID 6QBA) with binding surface in cyan (45). **(B)** Analysis of the affinity of POX9 (black) and its affinity-matured versions POX10 (blue) and POX11 (red). Enriched PXDN-binders were displayed on the surface of yeast, followed by incubation with several different concentrations of PXDN-con4, which was subsequently detected with anti-Penta-His-AF488 by flow cytometry. Average median fluorescence intensity (MFI) values ± SDs of three independent experiments are depicted. The lines represent a 1:1 binding model fitted to the data. **(C)** Surface plasmon resonance. Soluble, *E. coli* expressed sfGFP-POX10 fusion protein was immobilized on an SPR chip, followed by injection of various concentrations of PXDN-con4. Data (black lines) were fitted using a kinetic 1:1 binding model (red).

### PXDN protein expression in metastatic melanoma cell lines

To measure PXDN protein expression, we established an ELISA using the PXDN binding protein for capture and an in-house antibody for detection. We first confirmed that the PXDN binding protein recognized melanoma PXDN with a pulldown experiment using the medium of two invasive melanoma cell lines (NZM6 and NZM40, supplementary Figure S3A). SDS-PAGE and staining with Coomassie Brilliant Blue gave a band of an approximate molar mass of 500 kDa (theoretical non-glycosylated mass of trimeric full length PXDN is 490 kDa), which peptide analysis by MS identified as PXDN (Supplementary Figure S3B). The ELISA was first tested with NZM40, and gave a positive response with a linear dependence on the amount of lysate protein added (Supplementary Figure S3C). It was not possible to calibrate with PXDN-con4 because this shortened form does not contain the C-terminal region recognized by the antibody.

PXDN protein was detected in cell lysates from all the invasive metastatic melanoma cell lines (NZM6, NZM9, NZM11, NZM22 and NZM40), whereas it was scarcely detectable in the non-invasive cell lines (NZM12, NZM15, NZM42) and the normal melanocyte cell line HEMn-LP (Figure 3A). PXDN secretion to the medium was also detected in all of the invasive cell lines with much lower levels in the non-invasive ones (Figure 3B). The total amount of secreted PXDN in the medium was several times higher than in the cell lysate (Figure 3A & B). From these data we estimated approximate secretion rates of 1.5-5.8 times the intracellular content per 24 hours for all cell lines (Table 1). Next the highest expressing cell line NZM40 was exchanged into fresh culture medium and intracellular and secreted PXDN protein was measured over the next six (intracellular) and four (secreted) days (Figure 3C and 3D). PXDN protein was continuously secreted and increased significantly over time whereas the intracellular protein concentration stayed at a similar level. Significant positive correlations between mRNA expression and intracellular PXDN protein expression as well as PXDN protein secretion to the medium were seen for all eight melanoma cell lines as depicted in Figure 3E and F.

**Figure 3:**
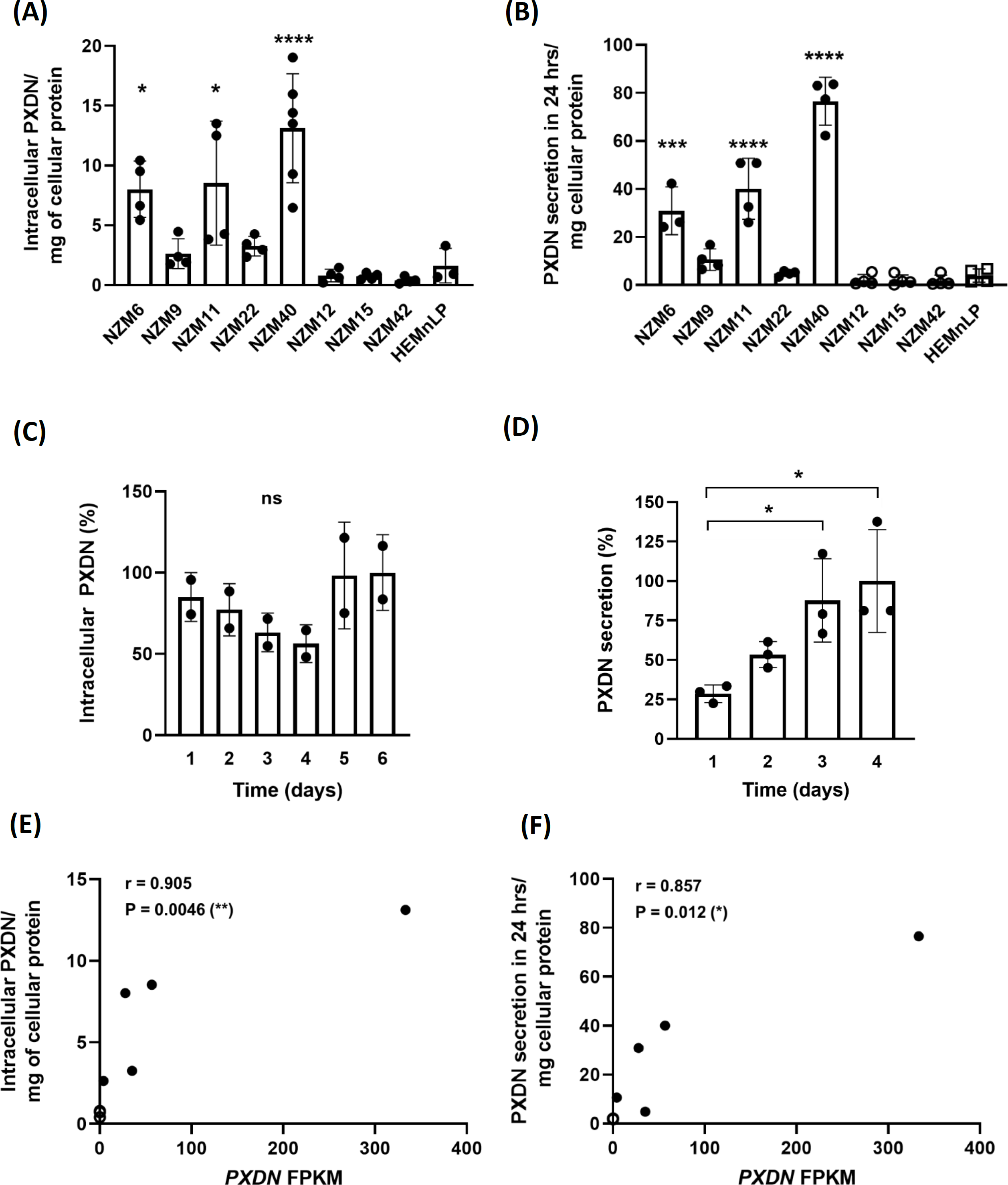
Relative levels of PXDN protein expression in melanoma cell lines determined by ELISA. PXDN protein expression in invasive (NZM6, NZM9, NZM11, NZM22, NZM40) and non-invasive (NZM12, NZM15, NZM42) cell lines and one normal melanocyte cell line HEMn-LP. **(A)** Intracellular PXDN determined by applying 50 μL of cell lysate per well (5 – 10 mg/mL protein) and signal was normalized to mg cellular protein. **(B)** PXDN secreted into the cell culture medium over 3 days measured by applying 50 μL from 10 mL medium to the ELISA. In (A) and (B) PXDN is expressed as arbitrary units based on absorbance in the ELISA. Bars represent the means of four biological repeats depicted as individual data points. *, *P* < 0.05, ***, *P* <0.001, ****, *P* <0.0001 by one-way ANOVA analysis followed by Tukey’s multiple comparisons between cell lines. **(C)** Intracellular PXDN levels of NZM40 measured over 6 days from 80 % confluence. Bars represent the means of two biological repeats, depicted as individual data points, performed in duplicate, 100 % refers to an absorbance of 0.5 at 405 nm. **(D)** Secretion of PXDN protein of NZM40 to the cell culture medium over time. Bars represent the mean value of two biological repeats shown as individual data points. *, *P* < 0.05 by one-way ANOVA analysis followed by Tukey’s multiple comparisons between measurements, 100 % refers to an absorbance of 0.4 at 405 nm. **(E)** Relationship between relative amounts of *PXDN* mRNA and cellular or **(F)** secreted protein expression of melanoma cell lines, analyzed by two-tailed Spearman correlation. PXDN protein units as for (A) and (B), (•) five invasive and (○) three non-invasive cell lines. The values for the three non-invasive cell lines overlap partly in (E) and fully in (F).

**Table 1:** Fold secretion of intracellular PXDN per 24 hours of melanoma cell lines and the melanocyte cell line HEMn-LP. Fold secretion was calculated from the amount of PXDN protein secreted to the medium in 24 hours relative to PXDN in the cell lysates estimated from the ELISA and MS data. Note NZM9 was not analyzed by MS and the non-invasive cell lines gave no signal in the MS assay.

| invasive | ELISA | MS | non-invasive | ELISA |
| --- | --- | --- | --- | --- |
| NZM6 | 3.9 | 3.9 | NZM12 | 2.6 |
| NZM9 | 4.0 | - | NZM15 | 2.8 |
| NZM11 | 4.7 | 3.4 | NZM42 | 4.3 |
| NZM22 | 1.5 | 2.0 | melanocyte | ELISA |
| NZM40 | 5.8 | 3.4 | HEMnLP | 2.4 |

We also used mass spectrometry to detect and quantify PXDN in the melanoma samples. The assay was based on tryptic digestion and analysis of two peptides with good signal strength identified from analysis of PXDN-con4 and confirmed by BLASTP search to be specific for PXDN. Good signal strength was obtained for both PXDN peptides in cell lysates as well as in the medium for the four high expressing cell lines examined (Figure 4A and 4B) and the relative values reflected those seen by ELISA. Values of ∼140 ng per mg cellular protein of cellular PXDN and ∼460 ng of PXDN secreted in 24 h were obtained for NZM40, with lower signals for the other investigated invasive cell lines (approximately 10-20 ng PXDN per mg of cellular protein and 20-80 ng PXDN secreted to the medium). No signal was obtained for the non-invasive cell lines (data not shown). Time course experiments with NZM40 (Figure 4C and 4D) confirmed those obtained by ELISA, showing a constant intracellular PXDN concentration and significant increase in the medium over time. These calculated values equate to low nanomolar concentrations of PXDN in the cell lysate of NZM40 (∼5 nM) and medium (∼1 nM) and respectively lower concentrations for the other cell lines. The MS results equated to a very similar rate of secretion of approximately 2-4 fold of the intracellular content of PXDN secreted per day (Table 1).

**Figure 4:**
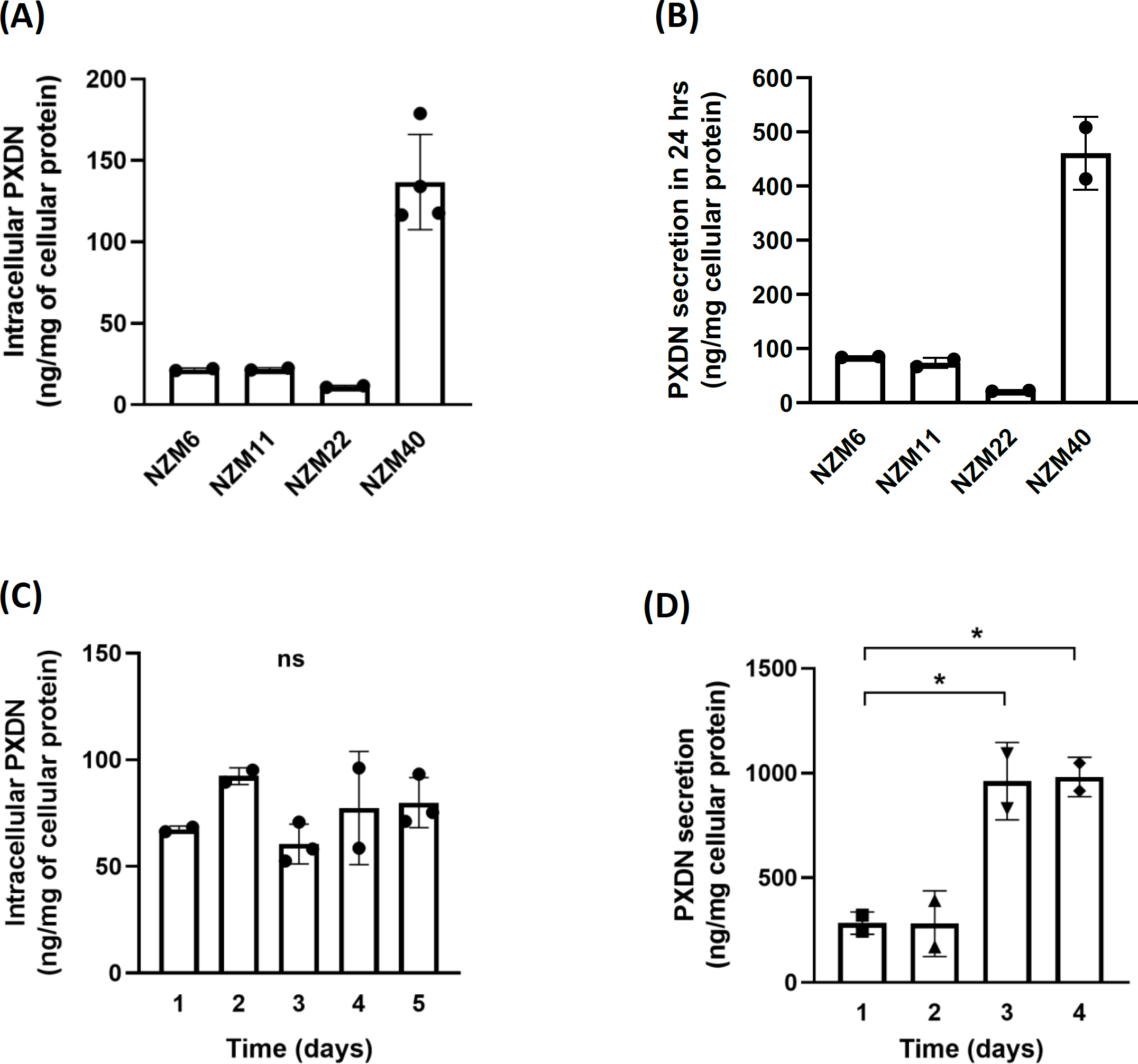
Levels of PXDN protein expression in invasive and non-invasive melanoma cell lines determined by MS. **(A)** Intracellular PXDN and **(B)** PXDN secreted over 24 hours was normalized to mg cellular protein. Bars represent the means of biological repeats, depicted as individual data. **(C)** Intracellular PXDN and **(D)** secreted PXDN measured over time in NZM40 cells. Bars represent the means of biological repeats, depicted as individual data points. Values are all normalized to mg cell protein and were estimated by comparing the peptide signal responses to those from a tryptic digest of a known amount of PXDN-con4 and assuming an equivalent MS response.

### Peroxidase activity of PXDN in metastatic melanoma cell lines

The ELISA was also used to measure PXDN activity with Amplex Red before the primary detection antibody was applied. A standard curve generated with PXDN-con4 displayed a concentration and time dependent increase of Amplex oxidation (Supplementary Figure S4A and S4B). It gave a threshold of ∼25 ng which required to be overcome before an increase in signal with PXDN-con4 concentration was detected. When cell lysates (250 – 500 µg cellular protein) or the medium from metastatic melanoma cell lines were applied to the ELISA, activity was only consistently detected in the highest PXDN expressing cell line NZM40 (Figure 5). Increasing amounts of NZM40 lysate protein gave a concentration dependence curve of cellular PXDN activity similar in shape to that observed for pure PXDN-con4, with the increase of PXDN activity becoming linear above a 50 µg threshold of total cellular protein (Supplementary Figure S4A and 4B). The response seen with 100-200 µg cell lysate protein was of the same order as that seen with 100-200 ng of PXDN-con4. Assuming a similar binding and activity of the two, this equates to ∼1 ng active PXDN per µg protein, somewhat higher than measured by MS. Confirmation that the assay was peroxidase-specific was obtained using phloroglucinol (PHG), a potent PXDN inhibitor, and azide, a general heme protein poison, both of which inhibited the enzymatic activity of NZM40 cell lysate as shown in Figure 5B. Active PXDN was consistently detected in the medium of NZM40 (Figure 5C) with the amount present increasing over time (Figure 5D). The signal for the NZM40 medium was much lower than that obtained for intracellular PXDN. This is most likely due to the high dilution factor of PXDN in the medium and the non-linearity of the activity signal at low concentrations, seen in both, the standard curve of PXDN-con4 (Supplementary Figure S4B) and in the loss of signal for activity at low concentrations of NZM40 cell lysate (Supplementary Figure S4C).

**Figure 5:**
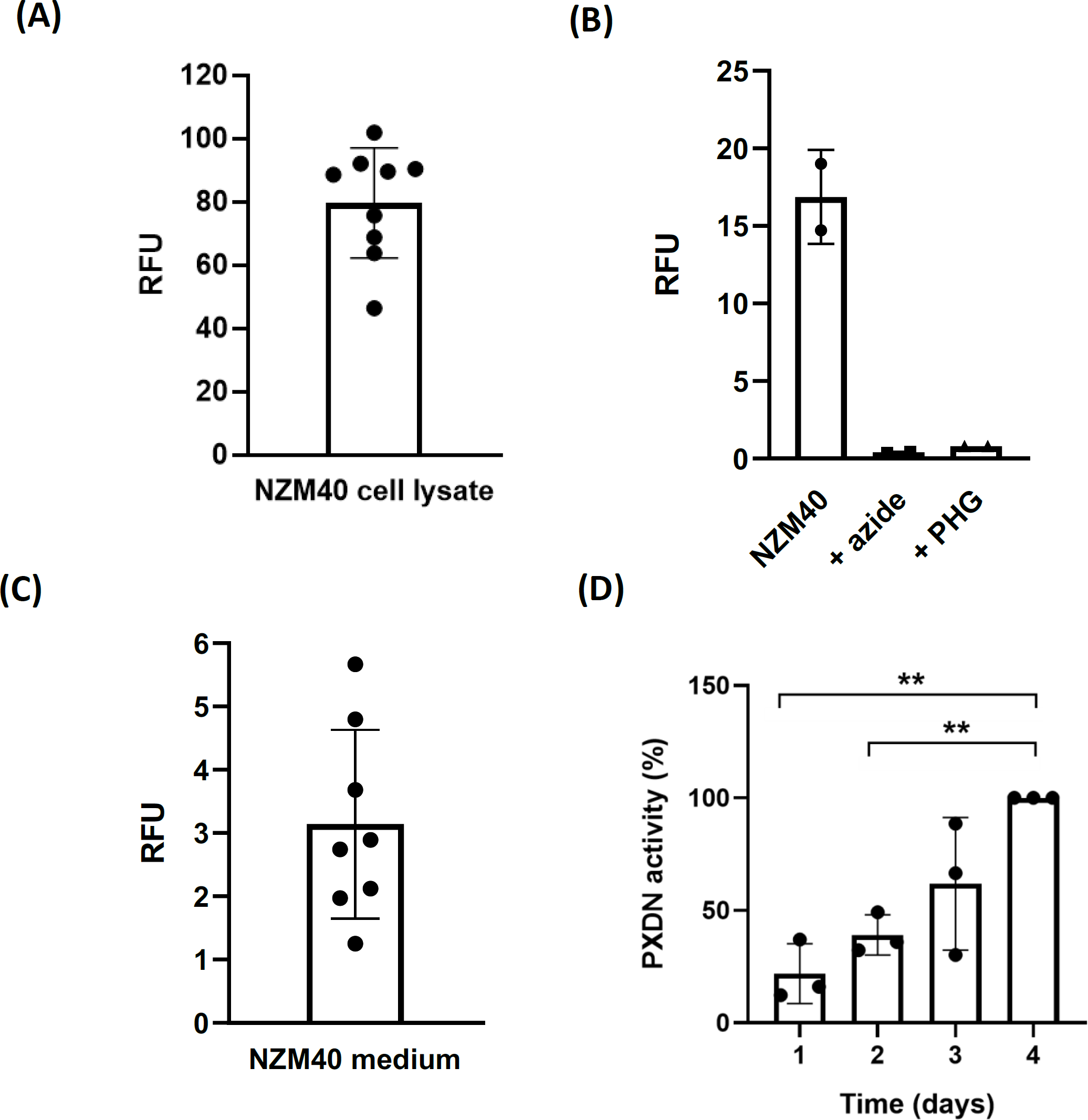
PXDN activity of NZM40 by ELISA. **(A)** PXDN activity detected in the cell lysate of NZM40 with cellular protein applied in the range of 250 – 500 μg. Bar represents mean ± SD of 9 independent experiments, each performed in duplicate. **(B)** Inhibition of PXDN activity of 100 μg of NZM40 cell lysate by the addition of 2 mM azide and 100 μM phloroglucinol (PHG), respectively. **(C)** PXDN activity detected in 50 μL of medium of NZM40 after three days of secretion. Bars represent mean ± SD of 8 independent experiments, each performed in triplicate. (D) Activity of PXDN detected in 50 μL of medium of NZM40 after each day for 4 days. Graph represents mean ± SD of three independent experiments performed in duplicate.

To establish the specificity of the ELISA for PXDN, both myeloperoxidase (MPO) (40 – 250 pg) and bovine lactoperoxidase (LPO) (80 – 500 pg) were analyzed. Neither gave a signal for activity or protein (n=2 independent experiments, data not shown). When used with a monoclonal antibody against MPO instead of the PXDN binding protein, 250 pg of MPO typically give a signal around 1000 RFU (46).

### Sulfilimine cross link formation in collagen IV by secreted melanoma PXDN

PXDN catalyzes the formation of sulfilimine cross links in the non-collagenous domain of collagen IV in ECM produced by epithelial cells. To establish whether PXDN secreted from melanoma cells can do this, we overlaid these cells on ECM from PFHR9 PXDN knockout cells. PHRF9 cells are differentiated, polarized mouse epithelial cells, often used to study ECM deposition. It has been shown that wild type PFHR9 cells form ECM with approximately 70 % collagen IV NCD dimers whereas PFHR9 PXDN knockout NCDs stay mainly uncross-linked and monomeric (< 15 % NCD dimers) (15, 38, 47). We first confirmed this for our system where PFHR9 WT and PXDN KO cells were grown for 8-10 days and NCD of collagen IV cross-linking was analyzed by SDS-PAGE (Figure 6A, lane 1 PFHR9 WT ECM and lane 2 PFHR9 PXDN KO ECM). The sulfilimine link of PXDN KO ECM was restored when PFHR9 WT cells were seeded confluently on top of PXDN knockout ECM (Figure 6A, lane 3). When the highest PXDN expressing melanoma cell line NZM40 was grown on PXDN KO ECM for 48 h, NCD cross link formation was restored to a similar degree (approx. 60 % dimer formation). Cross-linking was substantially decreased to approx. 10 % when PXDN was inhibited by PHG (Figure 6A, lane 4 and 5 respectively). The non-invasive, low PXDN expressing melanoma cell line NZM42 did not lead to cross link formation (Figure 6A, lane 6). Densitometry quantification is shown in Figure 6B. These results show that PXDN secreted by melanoma cells can cross-link collagen IV.

**Figure 6:**
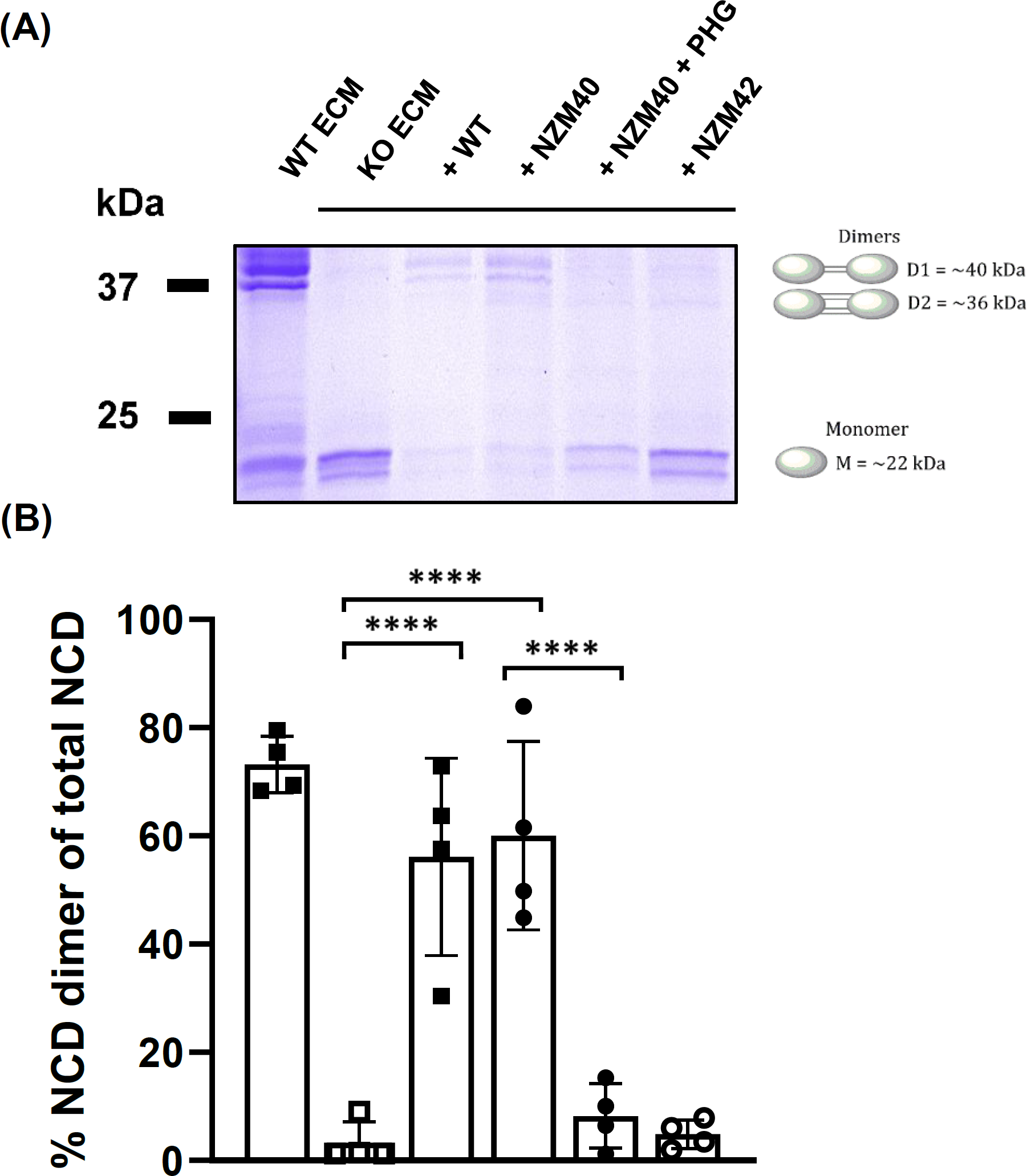
Sulfilimine cross link formation in the NCD of collagen IV by melanoma PXDN. **(A)** PFHR9 WT and PXDN KO cell lines were grown for 8 – 10 days to allow for extensive ECM deposition. Cells were hypotonically lysed to generate PFHR9 WT and PXDN KO ECM. PFHR9 WT cells, NZM40 and NZM42 were grown on top of the ECM isolated from PXDN KO cells for 24 or 48 hrs for PFHR9 WT and melanoma cells, respectively and sulfilimine link formation was analyzed by non-reducing SDS PAGE after collagenase digestion of collagen IV. PFHR9 WT ECM (lane 1), PFHR9 PXDN KO ECM (lane 2), KO ECM + WT cells (lane 3), KO ECM + NZM40 (lane 4), KO ECM + NZM40 + 100 µM PHG (lane 5) and KO ECM + NZM42 (lane 6). Representative gel of ≥ 4 independent biological repeat experiments. **(B)** Densitometry analysis of Coomassie Blue stained gels from four independent experiments as in (A) aligned under their respective gel lanes, mean values ± SD. The percentage of cross links was calculated as per the equation: % Dimer = (D1+D2×2)/(D1+D2×2+M) × 100, where D1 is the density of Dimer 1 (upper band) which contains one sulfilimine link and D2 is Dimer 2 (lower band) which contains two sulfilimine links, thus is multiplied by 2 to account for both bonds. ****, *P* <0.0001 by one-way ANOVA analysis followed by Tukey’s multiple comparisons.

### Peroxidasin activity leads to bromotyrosine formation in invasive metastatic melanoma cells

Sulfilimine formation in collagen IV implies PXDN-dependent HOBr formation. To test whether PXDN in melanoma cells forms HOBr, we measured the specific HOBr biomarker, 3-bromotyrosine, in cell protein from the highest PXDN expressing cell line, NZM40, and one low expressing cell line, NZM42 (Figure 7). Bromotyrosine was readily detectable in NZM40 cells and increased significantly when 100 μM or 200 μM bromide was present in the cell culture medium. The addition of PHG decreased the formation of bromotyrosine, confirming PXDN specificity. NZM42 gave only low levels of bromotyrosine which did not increase significantly when 200 μM bromide was added, further confirming PXDN-dependence of HOBr formation. These results demonstrate that PXDN in melanoma cells forms HOBr that can brominate tyrosine residues in proteins.

**Figure 7:**
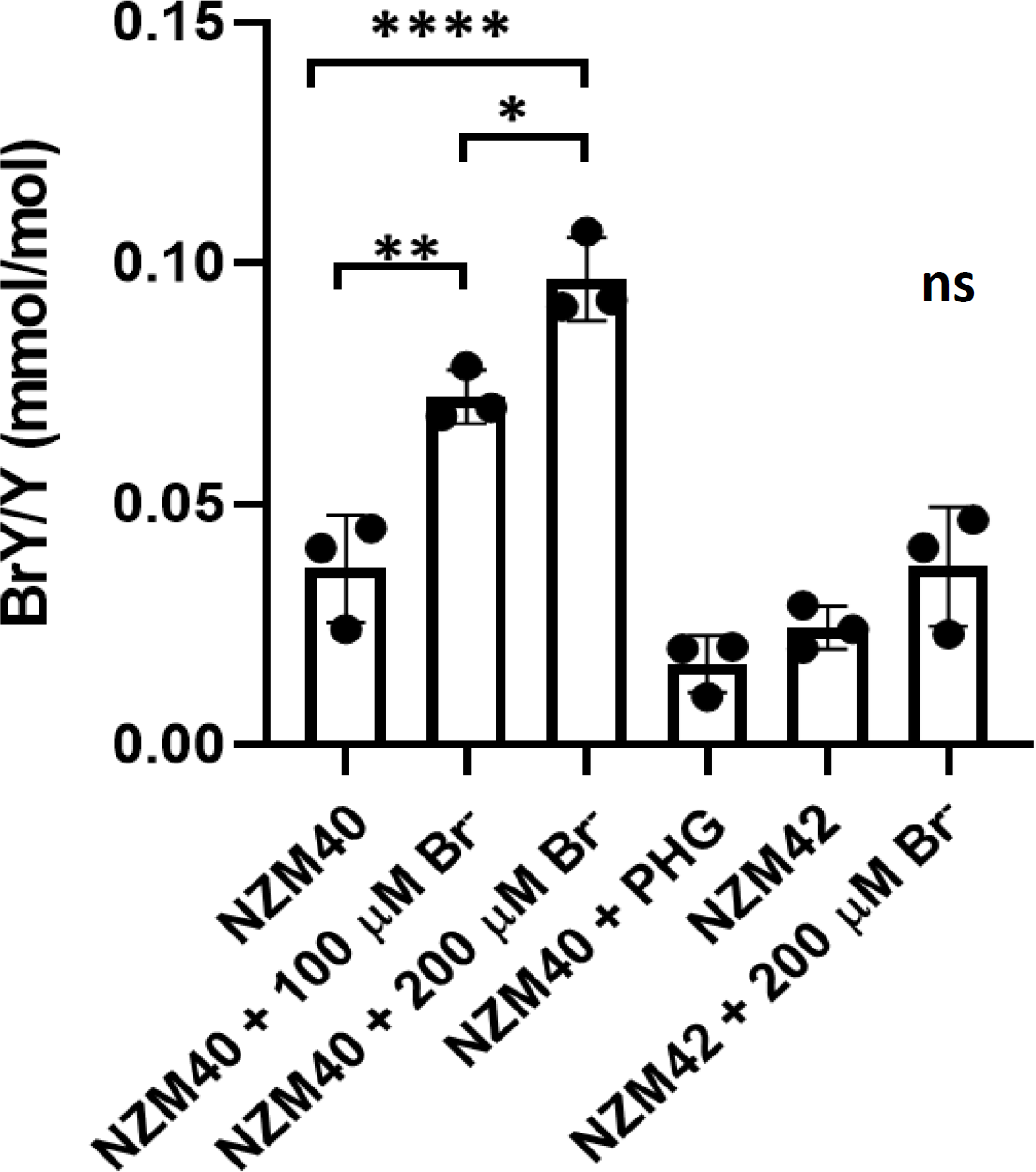
3-Bromotyrosine in melanoma cell lysates. 3-bromotyrosine was detected by MS in cell lysates of NZM40 and NZM42 and expressed relative to tyrosine content in hydrolyzed protein. Cells were grown for seven days and medium was exchanged daily and supplemented with 100 μM or 200 μM bromide or 100 μM PHG. Bars represent mean values ± SD of three independent experiments performed in triplicate. *, *P* < 0.05, **, *P* < 0.01, ****, *P* < 0.0001 by one-way ANOVA analysis followed by Tukey’s multiple comparisons.

## DISCUSSION

The work of Jayachandran and colleagues demonstrated high *PXDN* mRNA expression in invasive melanoma cell lines and identified PXDN as a determinant of melanoma cell invasion (2). We have extended these observations to a set of cell lines from the well characterized NZM metastatic melanoma panel to show on average 100 times higher *PXDN* mRNA expression in the five invasive cell lines when compared to the three non-invasive cell lines tested and a corresponding differential in expression of PXDN protein. In a related study of this panel of cells, *PXDN* was identified as one of the most differentially expressed genes (43). We also show that melanoma cell PXDN is secreted from the cells, it is active, and it generates HOBr which can form a specific cross link in collagen IV and undergo other reactions such as bromination of tyrosine residues in proteins.

In order to investigate PXDN protein expression we set up an ELISA using an in-house PXDN binding protein developed using a directed evolution yeast surface display approach (32, 34). Several binder families have been generated, recognizing at least two different epitopes on PXDN. This, plus their capability of being expressed as tagged forms or with a fluorescent readout, makes them a valuable tools for following native PXDN, which we are happy to make available. However, it should be noted that they do not work on denatured PXDN, e.g. with SDS-PAGE, as they recognize a small area of PXDN in its native 3D structure. Affinity matured POX10 was chosen as the most promising PXDN binder candidate for ELISA, based on a low *K*_d_ and absence of (or only low levels of) non-specific interactions as suggested by size exclusion chromatography. Binding to melanoma PXDN was demonstrated in a pulldown from the medium of PXDN-expressing cells and subsequent identification of the pulled down band by MS. Coomassie-stained SDS-PAGE showed very low non-specific binding and no cross-reactivity with MPO and LPO was seen in an activity assay. Further specificity to the ELISA is provided by the anti-PXDN antibody recognizing the C-terminal region (amino acids 1329 to 1479) which is unique for PXDN.

We detected PXDN protein by ELISA in the five invasive NZM cell lines, with very low levels in the non-invasive cell lines and in a melanocyte cell line, HEMn-LP. Relative levels were corroborated by MS analysis of PXDN peptides, and correlated well with the RNAseq values. The similarity of the non-invasive cells with HEMn-LP cells is consistent with their mRNA expression profile being similar to the melanocytic lineage (44). From the transcription data (Figure 1) a larger difference in protein expression levels between invasive and non-invasive cell lines might be expected. However, unknown mechanisms may regulate PXDN translation; for example, a specific microRNA (MiR-203a) was identified as a repressor of PXDN protein expression in a breast progenitor cell line (8).

PXDN in other cells has been shown to be a secreted protein that is translated in the endoplasmic reticulum and exported via the Golgi apparatus (14). This is also the case for melanoma cell PXDN. We found that intracellular levels measured over 6 days stayed constant but PXDN was continually released and accumulated in the cell culture medium. As calculated from the ELISA and MS data, approximately four times the intracellular PXDN content was secreted per day.

Coupling the ELISA with Amplex Red enabled detection of PXDN activity. It was able to detect a concentration-dependent increase in response with PXDN-con4. However, the assay showed a threshold limit of detection of ∼1 µg/mL (∼ 7 nM heme) which limited sensitivity. It gave a positive response with the highest expressing NZM40 cells and thus demonstrated that the intracellular PXDN was active, but gave no response with the other cell lines. The concentration dependence curve for NZM40 cell lysates also showed a lower limit threshold and it is most likely that PXDN is active in the cell lines with lower protein expression but levels are below the limit of detection in this assay. Activity measurements on the medium of NZM40 cells were very low due to dilution of secreted PXDN into a large volume, but they were significantly above baseline and increased over time, indicating that the secreted protein also has activity.

Whereas the ELISA protein assay provided data on relative content of PXDN in the different cell lines, it could not be calibrated with PXDN-con4 to give absolute levels. Comparing the MS response for common peptides from hydrolyzed protein from NZM40 lysate and PXDN-con4 gave a value of approximately 100 ng PXDN per mg cell protein. However, this value is dependent on the PXDN peptides giving similar responses in the two matrices and requires validation with an internal standard. Comparing the activities of NZM40 and PXDN-con4 suggests a higher content closer to 1000 ng/mg protein, but this value also has limitations as it assumes both have the same affinity for the binding protein and they are equally active. Further investigation is therefore needed to establish more definitive quantitative levels.

As cross-linking of collagen IV is the only identified function of mammalian PXDN, we wanted to know whether melanoma PXDN carries out this reaction. Studies with epithelial cells have shown that the cross link involves a sulfilimine bond between Met and hydroxylysine residues of two non-collagenous domain monomers and requires the generation of HOBr (48). Even though our transcriptome data show that melanoma cells secrete collagen IV, we found that the secreted ECM was insufficient for direct analysis of cross-linking. However, by growing melanoma cells on top of uncross-linked ECM deposited by a PXDN knockout mouse epithelial cell line (PFHR9), we were able to demonstrate that PXDN from NZM40 cells cross-linked collagen IV and restored NCD dimer formation to a similar level as that formed by WT PFHR9 cells (38). The specific inhibition of PXDN by PHG and the absence of detectable cross link formation with low-expressing NZM42 cells confirmed the requirement for PXDN. Even though hydrogen peroxide and bromide are required as PXDN substrates, neither was added to the cell culture medium in this experiment. This has also been observed with PFHR9 cells (37, 38) in which case traces of adventitious bromide in the medium were shown to be sufficient for collagen IV cross-linking (McCall 2014). As with PFHR9 cells(38), there must be sufficient endogenous generation of hydrogen peroxide for effective sulfilimine bond formation. Attempts to identify this source in epithelial cells have not yet been successful (49, 50) and further investigation is also needed to characterize the source of the hydrogen peroxide required for melanoma cell PXDN activity.

Our collagen IV cross-linking results clearly demonstrate that secreted melanoma cell PXDN is active and generates HOBr. HOBr is a highly reactive oxidant and one of the puzzling questions is whether formation of the highly specific sulfilimine cross link is concerted or is accompanied by other oxidative modifications to the ECM or other cell constituents. As the IgG motifs of PXDN are required for efficient sulfilimine formation (39) and the leucine-rich repeat domain facilitates direct binding to laminin (27), protein-protein interactions may lead to spatially directed release of HOBr and selectivity. However, as was found for PXDN in PFHR9 cells (47), we have shown that HOBr produced by melanoma cell PXDN during culture is not completely localized and some reacts with tyrosine residues on cellular protein to form 3-bromotyrosine. The level of tyrosine bromination is similar to that previously observed for cell-derived proteins from PFHR9 cells. In that study, bromination of ECM proteins was also measured and was about 10-fold higher than for cell proteins. This implies that bromination of extracellular proteins is also likely to be higher with the melanoma cells. As with collagen cross-linking, endogenous hydrogen peroxide was sufficient for tyrosine bromination in NZM40 cells, but the level was enhanced when bromide was added to the medium. This suggests some selectivity for cross-linking when bromide availability limits HOBr production.

Although 3-bromotryosine is a sensitive and stable marker for detecting HOBr, HOBr is much more reactive with other amino acids than tyrosine, particularly Cys, Met and Trp residues (19, 51). It also reacts with amino groups to form bromamines that although less reactive, are more selective for Cys and Met and can reach targets further from the source of generation. Thus, detection of 3-bromotyrosine indicates that other oxidative modifications to proteins and other biomolecules by HOBr are highly likely to have occurred. Other products formed by PXDN activity like HOSCN, which is highly thiol specific, and free radical species may cause additional oxidative modifications (18), including disulfides and dityrosine cross links, that are not fully reflected by 3-bromotyrosine. These modifications would be expected to affect proteins in the ECM where PXDN is secreted, and which is known for its low oxidative defense mechanisms(52). They should also occur in the endoplasmic reticulum prior to PXDN secretion and it is also possible that HOBr and longer-lived secondary oxidants could reach other compartments of the cell. Some of these reactions are potentially damaging, but others such as thiol oxidation could influence redox homeostasis and redox signaling pathways.

Summing up, we have demonstrated that high *PXDN* mRNA and protein expression is connected to the invasive phenotype in metastatic melanoma cell lines, corroborating previous evidence that it is a determinant of invasion. We have presented new evidence that (as established for high-expressing cells) melanoma PXDN is active as a peroxidase, it generates HOBr and cross-links collagen IV, and also causes other modifications to cell constituents such as tyrosine bromination. We also show that it is secreted and therefore likely to be present in the tumor microenvironment. A role for PXDN in the process of epithelial-mesenchymal transition under the control of the transcription factor Snail1 has previously been suggested (53). However, key questions are if and how its peroxidase activities are linked to invasiveness. One possibility is that there is a relationship with its specific action in forming sulfilimine linkages in collagen IV, potentially increasing the stiffness of the tumor matrix which is known to contribute to tumor progression and invasion (54). Alternatively, other reactions of HOBr, which we have shown to occur both intracellularly and extracellularly, could be relevant. Further characterization of the ECM and identification of specific HOBr-dependent modifications in PXDN-expressing cells and their relationship to invasive characteristics are needed before these questions can be answered and the role of high PXDN expression in tumor progression in melanoma as well as other cancers can be understood. If peroxidase activities of PXDN are shown to be detrimental, it may be possible to limit its activity by varying its substrates or the use of inhibitors thus making it a potentially attractive therapeutic target.

## ACKNOWLEDGEMENTS

This work was supported by the Marsden Fund of New Zealand (M. P., N. J. M., C. C. W.), the Canterbury Masonic Charitable Trust (M. P.), the Canterbury Medical Research Foundation (M. P.), the Austrian Science Fund (FWF Project W1224 – Doctoral Program on Biomolecular Technology of Proteins – BioToP) (B. S.), Maurice Wilkins Centre for Molecular Biodiscovery, Cancer Research Trust NZ, and HS and JC Anderson Trust (J. M., M. R. E.). We are grateful to Dr. Gautam Bhave from Vanderbilt University for the PFHR9 peroxidasin-knockout cell line.

## AUTHOR CONTRIBUTIONS

M. Paumann and C. C. Winterbourn designed the study with input from M. R. Eccles and J. Motwani on RNAseq and melanoma cell lines and M. W. Traxlmayr. J. Motwani provided the RNAseq data. M. Paumann designed, performed and analyzed ELISA, MS and bromotyrosine experiments with assistance of B. Bathish, L. Paton (MS method for PXDN detection), N. J. Magon (MS method bromotyrosine optimization). N. Kienzl developed and characterized the PXDN binding proteins and B. Sevcnikar developed the sfGFP binding protein, both under the mentorship of M. W. Traxlmayr, P. G. Furtmüller and C. Obinger. M. Paumann was responsible for writing the manuscript with input from all authors.

## Abbreviations

PXDN: human peroxidasin 1
PXDN-con4: human recombinant peroxidasin consisting of the four immunoglobulin domains and the peroxidase domain
Ig: immunoglobulin domain
POX: peroxidasin binding protein variant
ECM: extracellular matrix
BM: basement membrane
NCD: non-collagenous domain
MS: mass spectrometry
LC: liquid chromatography
MPO: myeloperoxidase
LPO: lactoperoxidase
PBS: phosphate buffered-saline
PHG: phloroglucinol
ERK1/2: extracellular signal-regulated kinases
AKT: protein kinase B
FAK: focal adhesion kinase
epPCR: error prone PCR
rcSso7d: reduced charge Sso7d
MFI: median fluorescence intensity
SPR: surface plasmon resonance

