## Supplementary Data for "Peroxidasin protein expression and enzymatic activity in metastatic melanoma cell lines are associated with invasive potential"

| ID | Q1 Mass | Q3 Mass | Collision energy (volts) |
|---|---|---|---|
| $^{675}$AGEIFER$^{681}$ y3$^{2+}$ | 411.4 | 451.3 | 23 |
| $^{675}$AGEIFER$^{681}$ y4$^{2+}$ | 411.4 | 564.3 | 23 |
| $^{675}$AGEIFER$^{681}$ b4$^{2+}$ | 411.4 | 371.2 | 23 |
| $^{675}$AGEIFER$^{681}$ y4$^{1+}$ | 821.4 | 564.3 | 65 |
| $^{1101}$AFFSPFR$^{1107}$ y5$^{2+}$ | 436.2 | 653.3 | 24 |
| $^{1101}$AFFSPFR$^{1107}$ b2$^{2+}$ | 436.2 | 219.1 | 24 |
| $^{1101}$AFFSPFR$^{1107}$ y4$^{2+}$ | 436.2 | 506.3 | 24 |

**Supplementary Table 1. Transitions of PXDN tryptic peptides monitored by multiple reaction monitoring LC-MS/MS.** All peptides were monitored and AGEIFER y3 and AFFSPFR y5 were used for quantification.

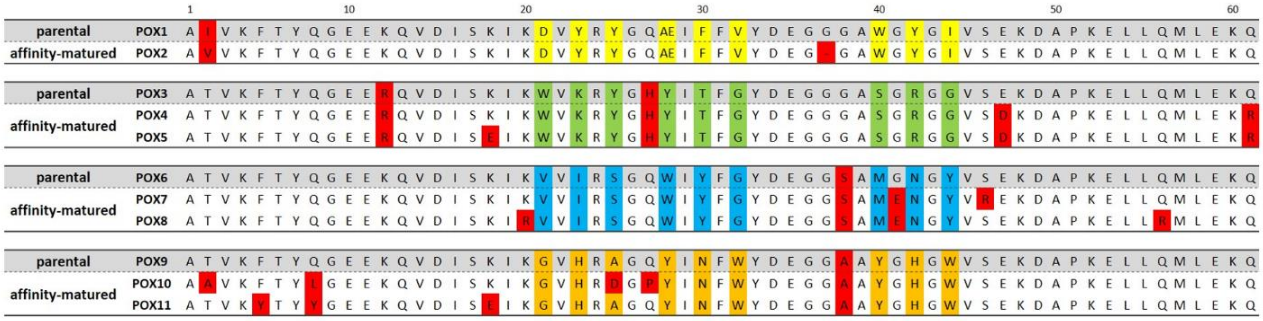


**Supplementary Figure S1: Sequences of enriched PXDN-specific binders.** Parental binders are marked in gray and their affinity-matured versions are shown below. The engineered binding surface (nine positions) is colored differently between the four sequence families (yellow, green, blue and orange, respectively). Point mutations introduced during error prone PCR are labeled in red. The first sequence family (POX1 and POX2) contains an insertion at position 28 and POX2 has a deletion at position 37.

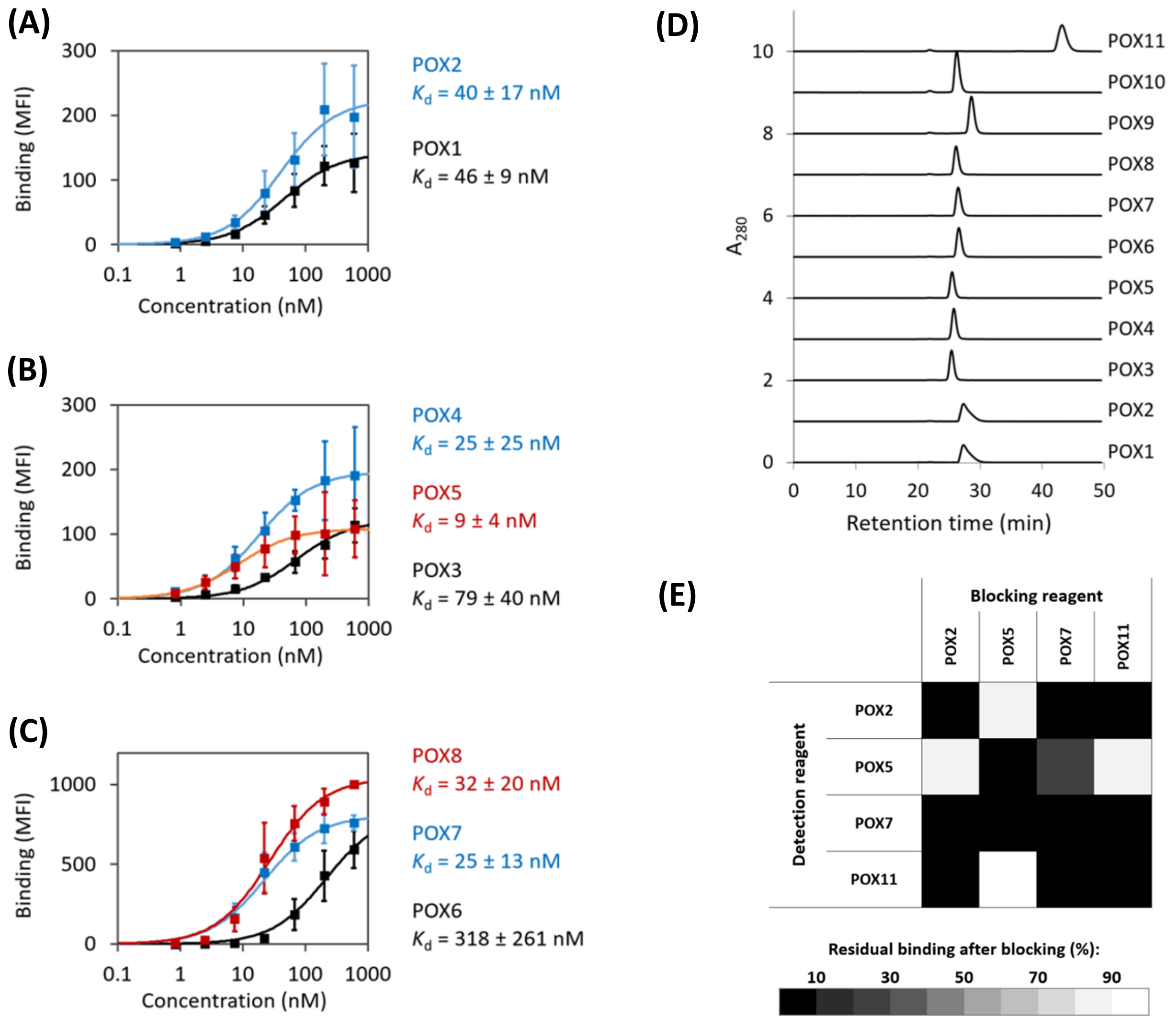


**Supplementary Figure S2: Detailed characterization of selected PXDN-binders. (A), (B)** and **(C)** Enriched PXDN-binders were displayed on the surface of yeast, followed by incubation with several different concentrations of PXDN-con4, which was subsequently detected with anti-Penta-His-AF488. Average MFI values ± SDs of three independent experiments are depicted. The lines represent a 1:1 binding model fitted to the data. Shown are three different sequence families based on POX1 (A), POX3 (B) and POX6 (C). Affinity-matured versions are depicted in red and blue. **(D)** Soluble binders were analyzed by size exclusion chromatography: 25 µg of the respective binder was injected onto a Superdex 200 10/300 GL column (GE Healthcare) using a Shimadzu prominence LC20 HPLC system equipped with a refractive index detector

(RID-10A, Shimadzu) and a diode array detector (SPD-M20A, Shimadzu). Samples were centrifuged and filtered before injection using 0.1 µm Ultrafree-MC filters (Merck Millipore; 3 min, 12,000 g, RT). Running buffer was PBS with 200 mM NaCl and the flow rate was 0.75 mL/min. Some mutants showed prolonged retention times during SEC, potentially caused by non-specific interaction with the column matrix. **(E)** Competition experiment. Binders POX2, POX5, POX7 and POX11 were displayed on the surface of yeast and tested for binding to PXDN-con4 (10 nM) that had been pre-incubated with soluble binders (1 µM, 10 min) as indicated. Equal volumes of pre-incubated binder and surface displayed binder were mixed. After 30 min, cells were centrifuged and washed with PBS supplemented with 0.1 % bovine serum albumin, followed by secondary staining with 1.7 µg/mL anti-HA-Alexa Fluor 647 and 2 µg/mL anti-Penta-His-Alexa Fluor 488 for detection of His-tagged PXDN-con4 bound to the yeast cells. The blocking capacity of the soluble binder was determined by comparison with PXDN-con4 binding in the absence of any soluble binder. Averages of two independent experiments are shown.

(A)

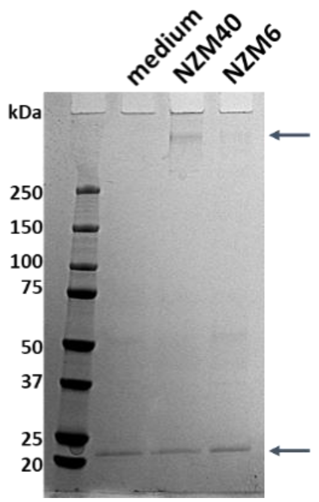


(C)

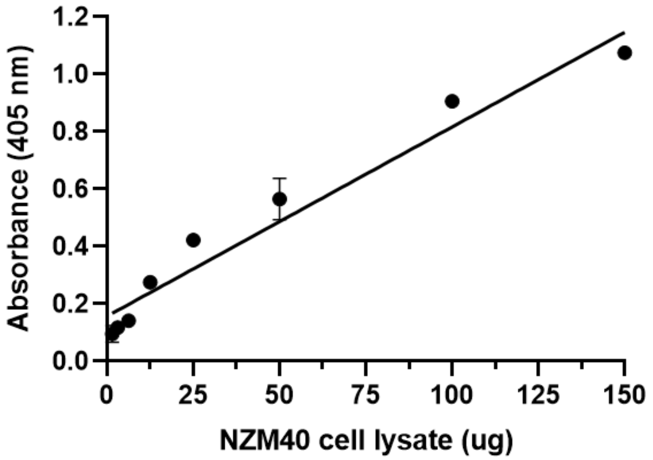


(B)

MAKRSRGPGRRCLLALVLFCAWGTLAVVAQKPGAGCPSRCLCFRTTVRCMHLLLEAVPAVAPQTSILDLRFNRIREIQPGAFRRLRNLNTLLLNNNQ
IKRIPSGAFEDLENLKYLYLYKNEIQSIDRQAFKGLASLEQLYLHFNQIETLDPDSFQHLPKLERLFLHNNRITHLVPGTFNHLESMKRLRLDSNTL
HCDCEILWLADLLKTYAESGNAQAAAICEYPRRIQGRSVATITPEELNCERPRITSEPQDADVTSGNTVYFTCRAEGNPKPEIIWLRNNNELSMKTD
SRLNLLDDGTLMIQNTQETDQGIYQCMAKNVAGEVKTQEVTLRYFGSPARPTFVIQPQNTEVLVGESVTLECSATGHPPPRISWTRGDRTPLPVDPR
VNITPSGGLYIQNVVQGDSGEYACSATNNIDSVHATAFIIVQALPQFTVTPQDRVVIEGQTVDFQCEAKGNPPPVIAWTKGGSQLSVDRRHLVLSSG
TLRISGVALHDQGQYECQAVNIIGSQKVVAHLTVQPRVTPVFASIPSDTTVEVGANVQLPCSSQGEPEPAITWNKDGVQVTESGKFHISPEGFLTIN
DVGPADAGRYECVARNTIGSASVSMVLSVNVPDVSRNGDPFVATSIVEAIATVDRAINSTRTHLFDSRPRSPNDLLALFRYPRDPYTVEQARAGEIF
ERTLQLIQEHVQHGLMVDLNGTSYHYNDLVSPQYLNLIANLSGCTAHRRVNNCSDMCFHQKYRTHDGTCNNLQHPMWGASLTAFERLLKSVYENGFN
TPRGINPHRLYNGHALPMPRLVSTTLIGTETVTPDEQFTHMLMQWGQFLDHDLDSTVVALSQARFSDGQHCSNVCSNDPPCFSVMIPPNDSRARSGA
RCMFFVRSSPVCGSGMTSLLMNSVYPREQINQLTSYIDASNVYGSTEHEARSIRDLASHRGLLRQGIVQRSGKPLLPFATGPPTECMRDENESPIPC
FLAGDHRANEQLGLTSMHTLWFREHNRIATELLKLNPHWDGDTIYYETRKIVGAEIQHITYQHWLPKILGEVGMRTLGEYHGYDPGINAGIFNAFAT
AAFRFGHTLVNPLLYRLDENFQPIAQDHLPLHKAFFSPFRIVNEGGIDPLLRGLFGVAGKMRVPSQLLNTELTERLFSMAHTVALDLAAINIQRGRD
HGIPPYHDYRVYCNLSAAHTFEDLKNEIKNPEIREKLKRLYGSTLNIDLFPALVVEDLVPGSRLGPTLMCLLSTQFKRLRDGDRLWYENPGVFSPAQ
LTQIKQTSLARILCDNADNITRVQSDVFRVAEFPHGYGSCDEIPRVDLRVWQDCCEDCRTRGQFNAFSYHFRGRRSLEFSYQEDKPTKKTRPRKIPS
VGRQGEHLSNSTSAFSTRSDASGTNDFREFVLEMQKTITDLRTQIKKLESRLSTTECVDAGGESHANNTKWKKDACTICECKDGQVTCFVEACPPAT
CAVPVNIPGACCPVCLQKRAEEKP

**Supplementary Figure S3. (A)** Pulldown of melanoma PXDN using the PXDN binding protein. The binding protein was immobilized in a high binding ELISA microplate followed by a blocking step as described for the ELISA. Medium collected from two PXDN expressing cell lines, NZM6 and NZM40, was applied to the plate (250 μL/well, 24 wells, 2 h at 37 °C). After three PBS washing steps 1x non-reducing SDS loading buffer was used to elute all proteins bound to the wells. Samples were resolved on a 4-20 % SDS PAGE and stained with Coomassie Brilliant Blue R250. Cell culture medium control in lane 1, NZM40 and NZM6 medium after 72 hours of PXDN secretion. Band at ~500 kDa was excised (top arrow) and identified as PXDN. Band at ~22 kDa PXDN binding protein ($His_6$-SUMO-POX10 fusion protein, bottom arrow). Representative gel of three independent experiments. **(B)** Identification of PXDN by MS. Excised gel band was digested with trypsin and analyzed by MS. Identified peptides are depicted in white highlighted in black and comprise an amino acid sequence coverage of 16 %. **(C)** Linear concentration dependence of

PXDN protein signal with increasing amount of NZM40 cell lysate applied to the ELISA. Representative data of three independent experiments performed in duplicate, mean ± SD.

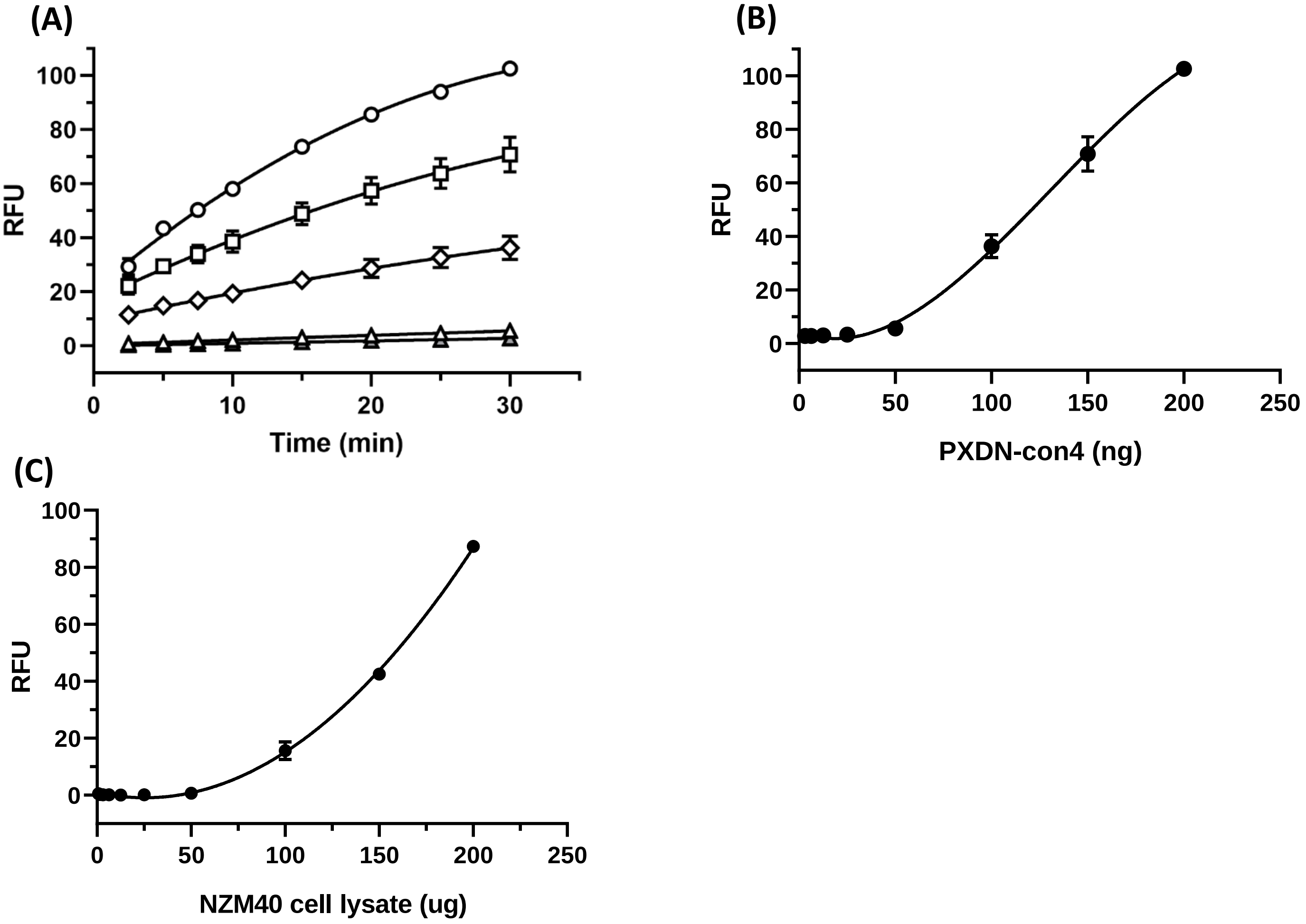


**Supplementary Figure S4. Oxidation of Amplex Red by PXDN-con4 and PXDN NZM40 cell lysate in ELISA.** **(A)** Concentration dependent increase of Amplex Red oxidation by PXDN-con4 over time (▲ blank, Δ 50 ng, ◊ 100 ng, □ 150 ng, ○ 200 ng). Representative graph of ≥ three independent experiments performed in duplicate displaying mean ± SD. **(B)** Standard curve created at 30 min of Amplex Red oxidation (3 – 200 ng of PXDN-con4). **(C)** Concentration dependence of PXDN activity of NZM40 cell lysate. Representative graph of three independent experiments, range ± SD of duplicate measurements.
